# Genome-scale metabolic modelling when changes in environmental conditions affect biomass composition

**DOI:** 10.1101/2020.12.03.409565

**Authors:** Christian Schulz, Tjaša Kumelj, Emil Karlsen, Eivind Almaas

**Affiliations:** Department of Biotechnology and Food Science, NTNU - Norwegian University of Science and Technology, Trondheim, Norway; K.G. Jebsen Center for Genetic Epidemiology Department of Public Health and General Practice, NTNU - Norwegian University of Science and Technology, Trondheim, Norway

## Abstract

Genome-scale metabolic modeling is an important tool in understanding metabolism, by enhancing collation of knowledge, interpretation of data, and prediction of metabolic capabilities. A central assumption in the construction and use of genome-scale models is that the *in vivo* organism is evolved for optimal growth, where growth is represented by flux through a biomass objective function (BOF). While the specific composition of the BOF is crucial, its formulation is often inherited from similar organisms due to the experimental challenges associated with its proper determination.

However, a cell’s macro-molecular composition is not fixed and it responds to changes in environmental conditions. As a consequence, initiatives for the high-fidelity determination of cellular biomass composition have been launched. Thus, there is a need for a mathematical and computational framework capable of using multiple measurements of cellular biomass composition in different environments. Here, we propose two different computational approaches for directly addressing this challenge: Biomass Trade-off Weighting (BTW) and Higher-dimensional-plane InterPolation (HIP).

In lieu of experimental data on biomass composition-variation in response to changing nutrient environment, we assess the properties of BTW and HIP using three hypothetical, yet biologically plausible, BOFs for the *Escherichia coli* genome-scale metabolic model *i*ML1515. We find that the BTW and HIP formulations have a significant impact on model performance and phenotypes. Furthermore, the BTW method generates larger growth rates in all environments when compared to HIP. Using acetate secretion and the respiratory quotient as proxies for phenotypic changes, we find marked differences between the methods as HIP generates BOFs more similar to a reference BOF than BTW. We conclude that the presented methods constitute a first conceptual step in developing genome-scale metabolic modelling approaches capable of addressing the inherent dependence of cellular biomass composition on nutrient environments.

**Author summary:** Changes in the environment promote changes in an organism’s metabolism. To achieve balanced growth states for near-optimal function, cells respond through metabolic rearrangements, which may influence the biosynthesis of metabolic precursors for building a cell’s molecular constituents. Therefore, it is necessary to take the dependence of biomass composition on environmental conditions into consideration. While measuring the biomass composition for some environments is possible, and should be done, it cannot be completed for all possible environments.

In this work, we propose two main approaches, BTW and HIP, for addressing the challenge of estimating biomass composition in response to environmental changes. We evaluate the phenotypic consequences of BTW and HIP by characterizing their effect on growth, secretion potential, respiratory efficiency, and gene essentiality of a cell.

Our work constitutes a first conceptual step in accounting for the influence of growth conditions on biomass composition, and in turn the biomass composition’s effect on metabolic phenotypic traits, within constraint-based modelling. As such, we believe it will improve the relevance of constraint-based methods in metabolic engineering and drug discovery, since the biosynthetic potential of microbes for generating industrially relevant products or drugs often is closely linked to their biomass composition.

## Introduction

The constraint-based reconstruction and analysis (COBRA) framework allows for the system-level analysis of genome-scale metabolism in microbes [1]. This framework has been used to construct metabolic models and knowledge bases for a large number of microorganisms with industrial and medical applications [2, 3]. Despite that the premise of COBRA is quite simple, the methodology is able to capture essential parts of the vast complexity of a full metabolism. In its simplest formulations, a COBRA approach such as flux balance analysis (FBA) is based on a set of linear equations corresponding to biochemical reactions, for which reasonable biological constraints are applied. This set of mathematical relations is subsequently converted into a linear program that is optimized with regards to a biologically plausible objective [1]. Typically, this objective is chosen to be the biomass objective function (BOF); a pseudo-reaction that utilizes the cellular metabolic network to consume nutrient resources. The output of the BOF is intended as a stoichiometrically balanced representation of the organism’s *in vivo* biomass composition for a given nutrient environment. The use of BOF as a cellular objective is a reasonable assumption, as an organism’s ability to quickly replicate is a property that often provides a fitness benefit [4, 5].

Given the widely accepted use of the BOF as an objective for genome-scale constraint-based metabolic modelling [3, 6, 7], relatively little attention has been given from this research field to exactly what the detailed composition of a cell is. In many cases, the formulation of a BOF is based on assumptions of similarity to related organisms; a chain of similarity that in many cases goes back to early model generations [8–16]. This has been an approach born out of necessity, as high-quality experimental determination of the detailed biomass composition of an organism is far from a trivial exercise [17, 18]. However, the detailed formulation of the BOF will affect phenotypic predictions [16, 19, 20].

The composition of the biomass closely reflects the metabolic state of the cell [7]. Changes in the environment will trigger changes in gene expression that eventually adjust or radically change the production of certain compounds, which over time will result in a different biomass composition [21]. In metabolism and macro-molecular expression models (ME-model), a stable structural composition is assumed where the (biomass) composition of proteins and transcripts are dependent on the nutrient environment [22–24], and these models typically contain tens of thousands of reactions. In contrast, the metabolic model (M-model, hereafter simply referred to as model) usually assumes a constant biomass composition. With typically just a few thousand reactions, this model type is significantly smaller and depend on accurate laboratory biomass component determination. These biomass components include precursor metabolites [25], DNA, RNA, proteins, lipids, coenzymes and cofactors, solutes, and more [7, 9, 20, 25–27]. During certain forms of starvation, the cell might also elect to accumulate some compounds for later use. An example of this is poly(3-hydroxybutyrate) (PHB) which can accumulate up to 87% of dry weight in *Alcaligenes latus* in a nitrogen limited environment [28].

Naturally, a range of important metabolic phenotypic traits change with biomass composition, such as growth potential, knock-out predictions, secretion rates, biosynthetic potential of industrially relevant products, or drug sensitivity [7, 16, 29, 30]. Based on this, one would also expect variations in nutrient environment to affect metabolic phenotypes due to their dependence on biomass composition. Therefore, measuring the biomass composition of an organism, not just once, but for a range of conditions relevant for the task at hand, has the potential for significantly improving modeling predictions.

However, measuring the biomass composition for every relevant combination of environmental parameters is unrealistic: There is a limit to the range of environmental or genetic conditions for which experimental measurements can be taken, which is an important reason for the popularity of GSMs. This raises the questions: How does one select among potentially multiple BOFs when modelling, and how can a limited number of experimentally determined BOFs be used to improve modelling predictions?

In this study, we propose two approaches to generate biomass compositions for a GSM that respond to changes in the nutrient environment based on linear combinations of available data. These methods, intended to be simple in both implementation and interpretation, are BTW (Biomass Tradeoff Weighting) and HIP (Higher-dimensional-plane InterPolation). They are based on the two assumptions that (i) the biomass composition depends on the nutrient environment, and (ii) similar environments yield similar biomass compositions. We explore the ramifications of BTW and HIP using the *Escherichia coli* model *i*ML1515 [20] with three fictional, yet biologically plausible, biomass compositions across varying glucose and ammonium uptake rates. Finally, we assess the impact of BTW and HIP on a set of key model characteristics, such as growth and acetate secretion potential, the respiratory state of the cell, and gene essentiality.

## Results and Discussion

The environment encoded in a GSM is solely incorporated in the bounds of the nutrient uptake rates. A typical minimal medium contains sources for carbon, nitrogen, phosphate, and sulphate, as well as some additional essential nutrients which depend on the organism. However, the particulars of a nutrient environment directly affects an organism’s macro-molecular composition [21], and we may thus consider the process of growth as a mapping between two sub-spaces: An environmental space where each uptake rate corresponds to a dimension, and the biomass space, where each biomass compound *BC_i_* corresponds to a dimension (see Materials and Methods for details). In this work, we limit our discussion to the case of only two uptake reactions, glucose and ammonium, for reasons of simplicity of presentation. Note however, the considerations presented in this work are also valid in higher-dimensional situations and are indeed motivated by such cases.

The glucose and ammonium uptake reactions generate a 2-dimensional environment space that represents all combinations of their flux uptake rates for the given ranges. We envision that for any given point in this 2-dimensional space, the biomass components of the cell would trend towards some ideal composition given enough time. This process would be guided by regulatory mechanisms responding directly to the metabolite concentrations and/or downstream consequences. Any point in this environment space should therefore have a corresponding point in biomass composition space. Note that, the mapping between the nutrient- and biomass component spaces is not necessarily a one-to-one mapping: Various environments could result in the same/similar biomass composition if, e.g the biomass composition is not monotonically dependent on one of the environment variables, and multiple points in the nutrient space could thus be mapped to a single point in the biomass component space. In lieu of high-quality data, however, we will make the simplifying assumption in the following discussion that the chosen uptake reactions (environmental dimensions) give rise to a 1-to-1 mapping between the two spaces.

In addition to metabolic uptake rates, other data sources relevant for any constraint-based modelling approach could be included as well. This includes experimentally determined growth rates or exchange rates of CO_2_ or acetate. All these data, as well as the BOF, are linked to a specific environment determined by the uptake reaction flux values.

In this work, we proceed by using a set of three artificial biomass compositions assumed to be measured for the same organism in three different environments: The first environment is characterized by nitrogen starvation (nitrogen limited - NL), the second by carbon starvation (carbon limited - CL), and the last is an environment where neither of these two elements are limiting (unlimited - UL). We will assume that the carbon source is glucose and the nitrogen source is ammonium, and the environments are specified by the uptake rates of the respective compounds (see Materials and Methods for details).

### Properties of the artificial BOFs for *i*ML1515

The BOF for the artificial nutrient-rich environment UL was based on the BOFs available in the *i*ML1515 model. The *core* BOF contains 31 compounds less than the *WT* BOF, and both contain the compound *adocbl*. In the standard minimal medium that is given in the model SBML-file, this compound prevents growth. Since a detailed curation of the model is beyond the scope of this work, we solved the discrepancy by simply removing this compound. We combined the two BOFs by taking the arithmetic mean of their respective biomass coefficients, and thereafter, scaled the resulting BOF to 1 g/gCDW^−1^. This step was implemented to assure a reliable generic BOF to imitate an environment with high availability of glucose and ammonium. Starting from this BOF, we generated the two artificial BOFs for nitrogen and carbon limited environments.

Briefly, we sorted the BOF compounds into groups, such as DNA, RNA, and protein. For the limited environments CL and NL, the relative amounts of the compounds in these groups were all scaled in a biologically inspired manner: For instance, we increased the relative fraction of compounds in the *carbohydrate* group in the nitrogen limited NL environment, and reduced it in the carbon limited CL environment. The details of the scaling are provided in Supplementary S1 Table and in Tab. 2 in the Materials and Methods section.

The three environments as defined by the specific maximum uptake flux rates for carbon and nitrogen are detailed in Tab. 1. The CL environment emulates a low uptake rate of glucose, while the NL environment emulates a low uptake rate of ammonia. The uptake rates shown in Tab. 1 are the upper bounds, with corresponding excretion rates being unconstrained. Note that, the BOFs and environments are created with the sole purpose of evaluating the methods for constraint-based simulations with multiple BOFs. Therefore, the defined uptake rates were chosen to be within the capabilities of a genome-scale model, not chosen to mirror physiolgical capabilities of *E. coli* in detail.

**Table 1.**
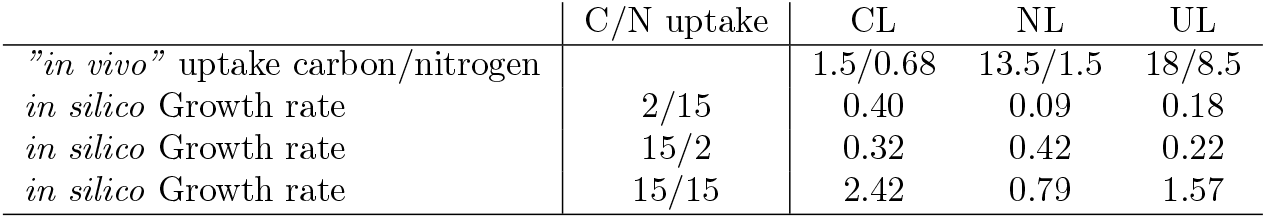
The artificial BOFs used in this model. CL is the carbon limited biomass, NL nitrogen limitation, and UL is the unlimited environment. The assumed maximum uptake flux rates are detailed for the respective environments. The growth rates are given in units of h^−1^, while the flux bound on the uptake rates are given in mmol gCDW^−1^ h^−1^.

|  | C/N uptake | CL | NL | UL |
| --- | --- | --- | --- | --- |
| "*in vivo*" uptake carbon/nitrogen |  | 1.5/0.68 | 13.5/1.5 | 18/8.5 |
| *in silico* Growth rate | 2/15 | 0.40 | 0.09 | 0.18 |
| *in silico* Growth rate | 15/2 | 0.32 | 0.42 | 0.22 |
| *in silico* Growth rate | 15/15 | 2.42 | 0.79 | 1.57 |

The CL and NL BOFs provide the largest growth rates within their respective environment when compared with each other. However, the UL BOF outperforms only the NL BOF in a rich environment. In many organisms, carbon (glucose) is the growth limiting compound, and the carbon-starvation biomass composition is created to survive with as little carbon as possible. Consequently, in an unconstrained environment, this BOF can generate more flux with the same carbon uptake rate, thus outperforming the UL BOF, as demonstrated in Fig. 1 panels **A** to **C**. If more biologically plausible constraints were imposed on the uptake reactions based on the available biomass composition, we are of the opinion that the CL biomass would not perform as well.

**Fig 1.**
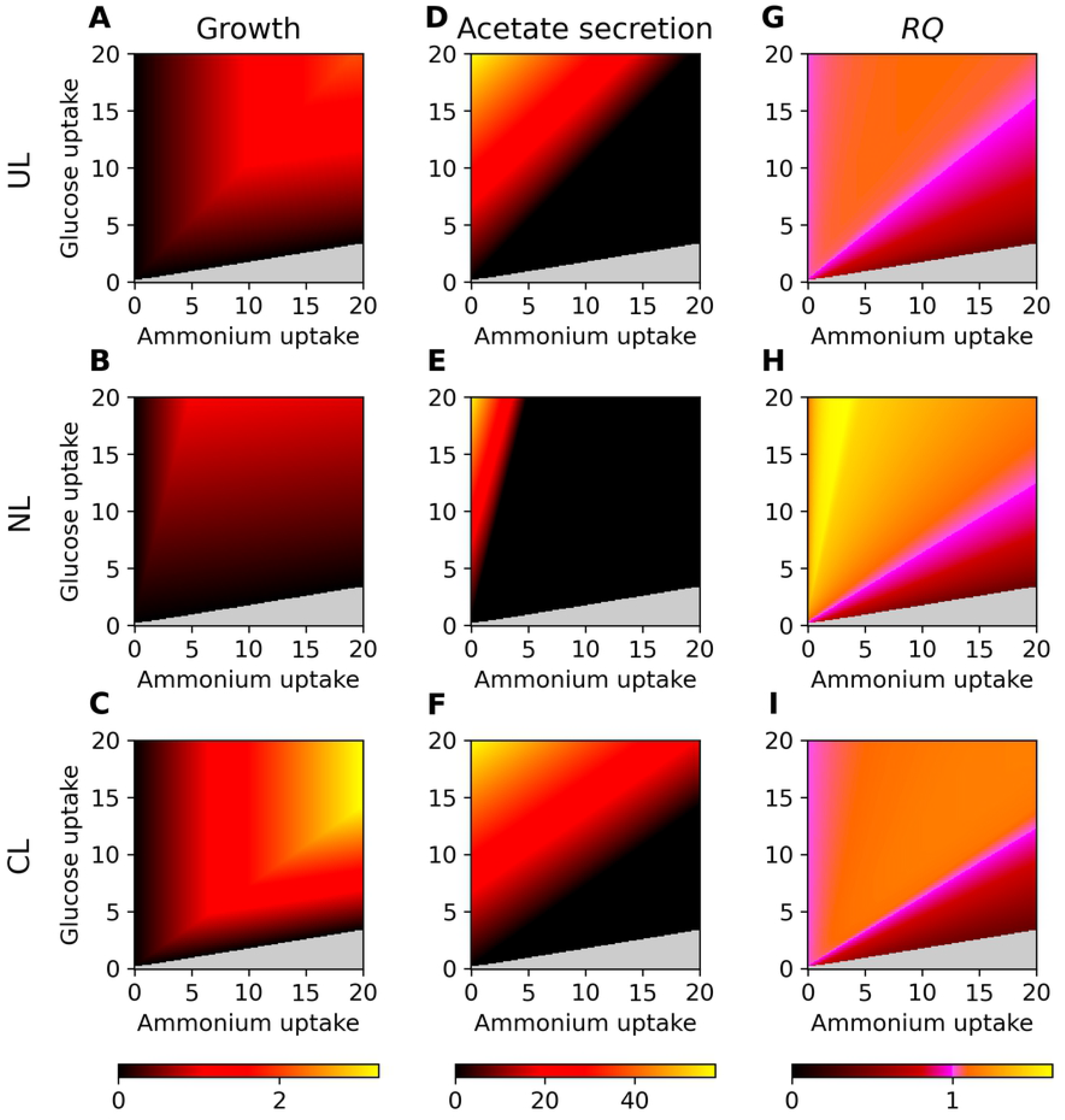
Comparison of the phenotypic properties of the three BOFs. In all panels, we vary the (fixed) glucose and ammonium uptake rates in units of mmol gCDW^−1^ h^−1^ along the horizontal and vertical axes, respectively. Non-growth areas are shown in light grey, and relative values of unity are represented by pink. Panels **A** to **C**, show the growth phenotype phase-planes with coloring according to growth rate. Maximal acetate secretion rates are shown in panels **D** to **F**. In panels **G** to **I**, we plot the respiratory quotient *RQ*.

Next, we explore the phenotypic properties of the three different BOFs. This is important to understand the impact of their formulation on the model and its phenotype predictions. To that end, we display key metabolic traits of the generated BOFs in Fig. 1. Panels **A** to **C** show the 2-D phenotype phase-planes of glucose against ammonium, where the calculated growth rate is represented by the color code. Note that all other uptake rates of the specified environment are unlimited. Furthermore, we draw the attention to the fact that the shown phenotype phase-planes are for the uptake of glucose plotted against ammonium, and should not be mistaken for the common plots of oxygen versus carbon source. We highlight that we only scaled the BOF compound coefficients; no compounds were added or removed, which is something one would expect to observe in an experimental BOF determination.

Not surprisingly, these panels show that the biomass composition affects the growth potential of cells in response to environmental changes. The CL biomass composition (with a reduced fraction of carbohydrates) generates a high growth potential in nitrogen and carbon rich environments, shown in panel **C**. In fact, the growth potential in a nutrient rich environment is even higher than the UL biomass, comparing panels **A** and **C**. This contrasts what one might expect to observe experimentally, as the UL BOF is constructed for a nutrient rich environment, whereas both limited BOFs are constructed for starving environments. However, as the ratio of carbon to nitrogen is CL < UL < NL, the CL BOF outperforms the other BOFs in the same environment, due to the fact that the same amount of carbon generates more flux through the BOF.

Acetate is the only byproduct that is secreted in both phenotypic states (respiration and fermentation), although the abundance of acetate secretion is higher in the fermentation state than in the respiratory state [25, 31]. Thus, the production of acetate is maximal at low oxygen availability, or low concentrations of other respiratory electron-chain acceptors, but non-zero in the presence of oxygen [32]. Fig. 1 panels **D** to **F** shows the secretion potential profile of acetate with respect to the biomass composition. In these calculations, we have maximized acetate production subject to optimal growth constraint. Large parts of the profile are in black, in areas of limited acetate production. This indicates the metabolism’s tendency towards a respiratory state. Note that, the maximal acetate production is in areas of a low nitrogen uptake or a high C/N ratio. Especially for the NL BOF, the acetate production phase is small, however, the phase transition gradient is very sharp. Acetate secretion is associated with lower growth [25, 31]. The model shows this behaviour in Fig. 1 panels **A** to **F**, where low nitrogen-carbon rich environments generate higher acetate secretion, which corresponds to low growth rates.

The respiration potential is measured by the respiratory quotient *RQ* and shown in Fig. 1 panels **G** to **I**. *RQ* is defined as the ratio between secreted carbon dioxide and absorbed oxygen *v*(CO_2_)/*v*(O_2_). To reasonably calculate RQ, we perform a multi-level optimization: First, we maximize growth using the respective BOF. Second, we maximize for acetate secretion subject to growth fixed at the maximal value. Finally, using a parsimonious FBA implementation, we minimize all reaction fluxes, including oxygen uptake and CO_2_ excretion, while keeping the BOF and the acetate fluxes at their previously determined (maximal) values.

*RQ* is another central phenotypic descriptor that one would expect it to be profoundly affected by the biomass composition, as it is connected to the the respiratory state of the cell [33]. As is evident from Fig. 1, the dependence of the *RQ* on the environmental parameters changes drastically with the different biomass compositions: *RQ* ≈ 1 is associated with fully oxidative respiratory metabolism, which is indicated by pink color in Fig. 1 panels **G** to **I**. In the respiratory state, the cell is able to fully oxidize glucose and produce flux trough the TCA cycle and electron transport chain [25, 34]. *RQ* < 1 represents fermentation, where pyruvate, as CO_2_ producing precursor of acetate formation, is excreted *in vivo*. In contrast, an *RQ* > 1 represents oxidative fermentation, where glycolytic activity with redox NAD^+^/NADH potential is enhanced and oxidative phosphorylation is impaired with the underlying decoupling of glycolysis and TCA cycle from oxidative phosphorylation [34]. This metabolic state is generally referred to as “overflow metabolism” [33]. All three metabolic states can be seen in the plots. Especially the NL BOF generates large areas where *RQ* > 1 for low ammonium uptakes; areas for which the CL and UL BOFs have an *RQ* ≈ 1. The area for *RQ* < 1, indicating a fermentative metabolism, is similar for the NL and CL BOFs, and larger for the UL BOF.

In sum, Fig. 1 shows how the chosen BOF formulations impact the performance of the *i*ML1515 genome-scale metabolic model using standard flux-balance analysis. This knowledge leads us to the main focus of this work: Given the availability of multiple BOF formulations, how may we use this knowledge in a flux-balance analysis formulation? Since it is only reasonable to assume that one may perform high-fidelity determination of a BOF for a limited set of conditions, we believe it necessary to develop heuristics that are capable of bridging this knowledge gap.

### Approaches to leverage multiple biomass composition data

Consider a situation where multiple BOFs are provided, and they are presumed to be clearly connected with (measured) environmental information. Each BOF is associated with a mapping from an *n*-dimensional environmental space onto an *m*-dimensional biomass component space. Here *n* corresponds to the number of relevant environmental parameters, and for reasons of simplicity, we will explore the case of *n* = 2 using carbon and nitrogen uptake as the environmental variables. The *m* dimensions correspond to the different possible molecular components in the BOF. The respective BOFs are constructed for different points in the environmental space, and we imagine them as resulting from measurements during those conditions in the wet lab. The challenge is then: If the metabolic behavior is of interest at locations in environment space that are between experimentally determined points, how does one combine the existing knowledge to infer relevant BOF composition?

In the following, we propose the two approaches of Biomass Tradoff Weighting (BTW) and Higher-dimensional-plane InterPolation (HIP). While they are both applicable to any value of *n*, we provide a visual presentation of their guiding principle in Fig. 2 using *n* = 1 for simplicity. Further details are provided in the Materials and Methods section. Simply put, the BTW approach allows a linear optimization algorithm to select the optimal combination of relevant (available) BOFs by setting their weights to unity in the objective vector **c** (Fig. 2**A**). The HIP algorithm (Fig. 2 **B**) interpolates between BOF compounds in biomass composition space by spanning the experimentally measured points with a linear plane. Additionally, in order to demonstrate an example of how these methods can be altered for greater utility and/or realism, we introduce an extended version of HIP, the Higher-dimensional-plane InterPolation-Iteration (HIP-I). HIP-I generates a BOF for a given environment using the HIP algorithm, after which the model is optimized without fixed uptake rates. The optimization step will oftentimes result in an optimal set of uptake flux rates that are not consistent with the coordinates for the BOF composition that was used. The new set of uptake rates is subsequently used to query the HIP algorithm for a new BOF. This procedure is iterated until it converges on a self-consistent point in nutrient and environment space.

**Fig 2.**
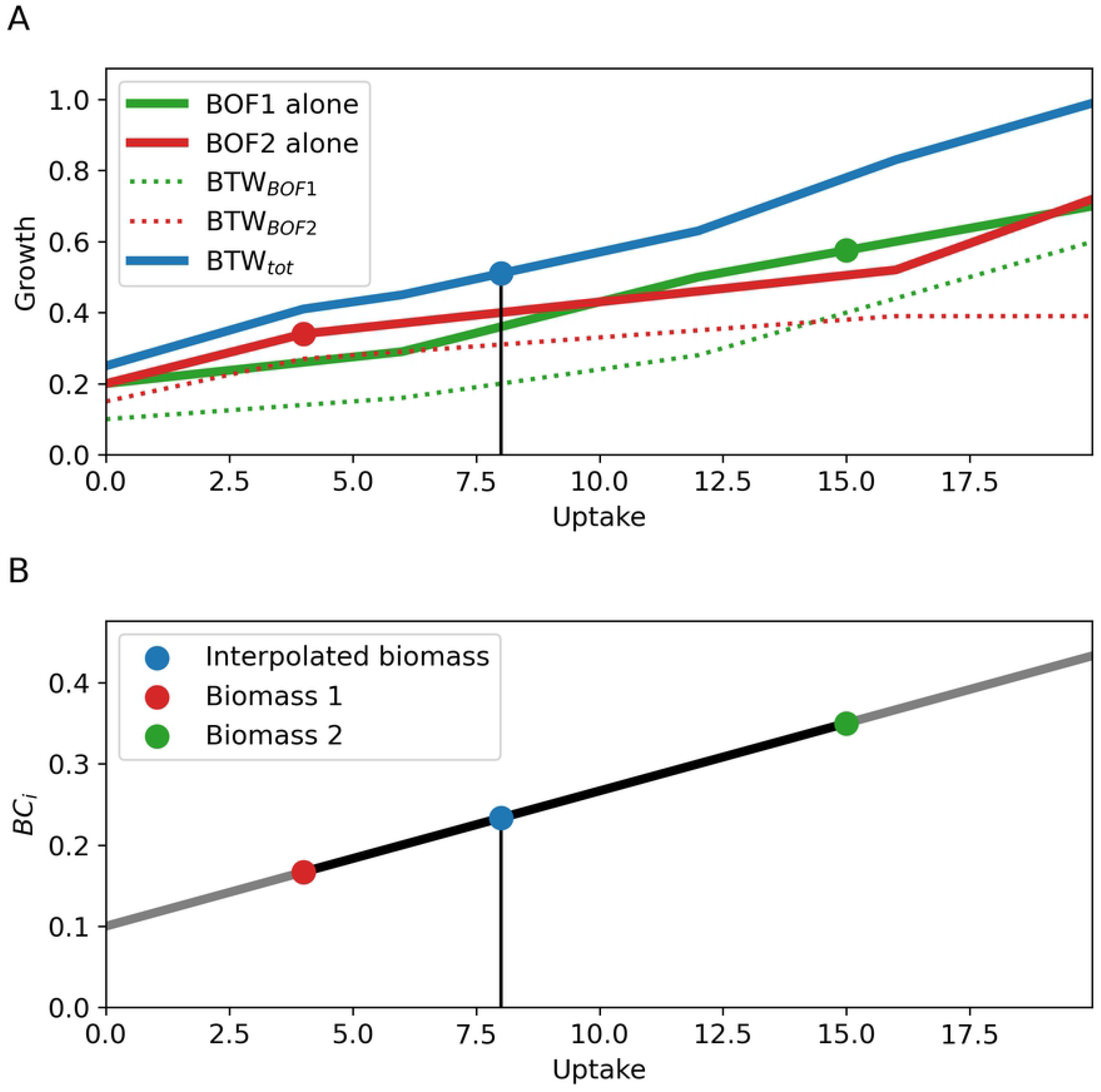
A simple illustration of the principles behind the Biomass Tradeoff Weighting (BTW) and Higher-dimensional-plane InterPolation (HIP) approaches. The green and red point represent a laboratory determined BOF (panel **A**) or the coefficient of biomass component *BC_j_* (panel **B**) for the given uptake. The blue point mark in both panels the requested environment defined by the respective uptake. **A**) An illustration of the BTW, where the linear optimization algorithm is allowed to select an optimal tradeoff between multiple BOFs in order to maximize total biomass production. The fluxes trough the different BOFs are combined for an optimal objective value, represented as the blue combined biomass. The green and red solid line represent the use of each single BOF, the dotted lines their contribution to the BTW BOF. **B**) A schematic depiction of HIP, which uses environmental parameters to interpolate between different known levels of biomass component coefficients *BC_i_* to generate coefficients for a new BOF. In contrast to BTW, the BOF is depending on the environment and nutrient data.

Since the genome-scale models of interest for computational analyses and biotechnological applications consist of a few to tens of thousands of reactions [23, 24], it is important to assess the computational efficiency of the proposed methods. The BTW approach only requires the time to add multiple biomasses once, and then setting each of their coefficients in the objective vector **c** to 1, also once. The model can be tested for any range of environments. The HIP approach requires the generation of a new BOF for the query point in nutrient space, which in our tests for the *i* ML1515 model and the two-dimensional environment space mentioned above, increased the mean compute time relative to just solving a standard FBA problem in the COBRA Toolbox, by 28.6% ± 1.2% over 10^4^ runs. Here, we have used the standard COBRA Toolbox functions for adding a reaction and changing the objective function. The time used for the model optimization itself does not change measurably in either of the two approaches.

Another point to consider is the “data hunger” of the different methods: BTW works with as little as one BOF (in the limit of one BOF, it reduces to regular single-BOF FBA), whereas interpolation requires measurements that span relevant environmental parameters. Interpolation is also applied assuming local linearity, and the reference mapping from environment to biomass composition should therefore have a certain level of resolution (in terms of the density of measurement points) for the assumption of linearity to hold.

For the HIP method, one could also envision non-linear interpolation functions between the experimentally determined BOF-points in biomass component space, but fitting these would require both a higher density of data points and additional biological insight into the mapping between environment-biomass component spaces. Consequently, the assumption of linearity is the simplest, and therefore the most prudent, given our current knowledge and lack of high-resolution biomass-composition data available.

While the HIP method is designed to interpolate between measurement points, its formulation also allows the extrapolation outside. Such extrapolation might in many cases be reasonable, for example close proximity to the measurement points, but extra care must then be taken. The evidence-base becomes shakier and extrapolation past a line drawn from a positive coefficient to a zero will result in a negative coefficient. This equates a forced infusion of a given biomass component into the metabolism, coupled to growth. In such cases, this phenomenon should be handled. For some compounds, such as glycogen or PHB, it is expected that they would be present in infinitesimal amounts in a given environment: For example, a carbon storage compound such as glycogen will not be formed in appreciable amounts during carbon starvation. Other compounds, say for example alanine, are essential for an organism and should not become zero. Consequently, in order to extrapolate to a BOF outside the region spanned by the experimentally determined biomass coordinates, one must classify such compounds and include a scale for returning a non-negative coefficient. Here, for simplicity, we used 1% of the respective UL coefficient, though alternative approaches are possible, and indeed highly required. In the following, we explore the phenotypic consequences of BTW and HIP/HIP-I when using the set of three artificial BOFs as input for the approaches.

### Comparing the multiple-biomass analysis methods

While determining one method as being strictly “better” than another is not meaningful without high-resolution empirical data, some comparisons are still in order, and we conducted a direct comparison for the three methods. In the standard phenotype phase-plane mapping, one uses fixed values for the tested uptake fluxes, which is not commensurable with the HIP-I iterative framework. Thus, we may only directly compare BTW and HIP. However, given the way HIP-I is defined as an iterative process, its zero’th iteration is identical to HIP. Also, environmental uptake flux values that correspond to the self-consistent HIP-I solutions (fixed points) are identical to the HIP results in these points.

In Fig. 3, we calculate several phenotype phase-planes by varying the two environmental variables of glucose and ammonium (using the same approach and values as in Fig. 1), and plot these relative to the values presented in the UL column of Fig. 1. The absolute plots for HIP and BTW can be found in S3 Fig.1. The phenotype profiles shown are growth (panels **A**-**B**), growth relative to UL (**C**-**D**), acetate secretion relative to UL (**E**-**F**) and respiratory quotient *RQ* relative to UL (panels **G**-**H**).

**Fig 3.**
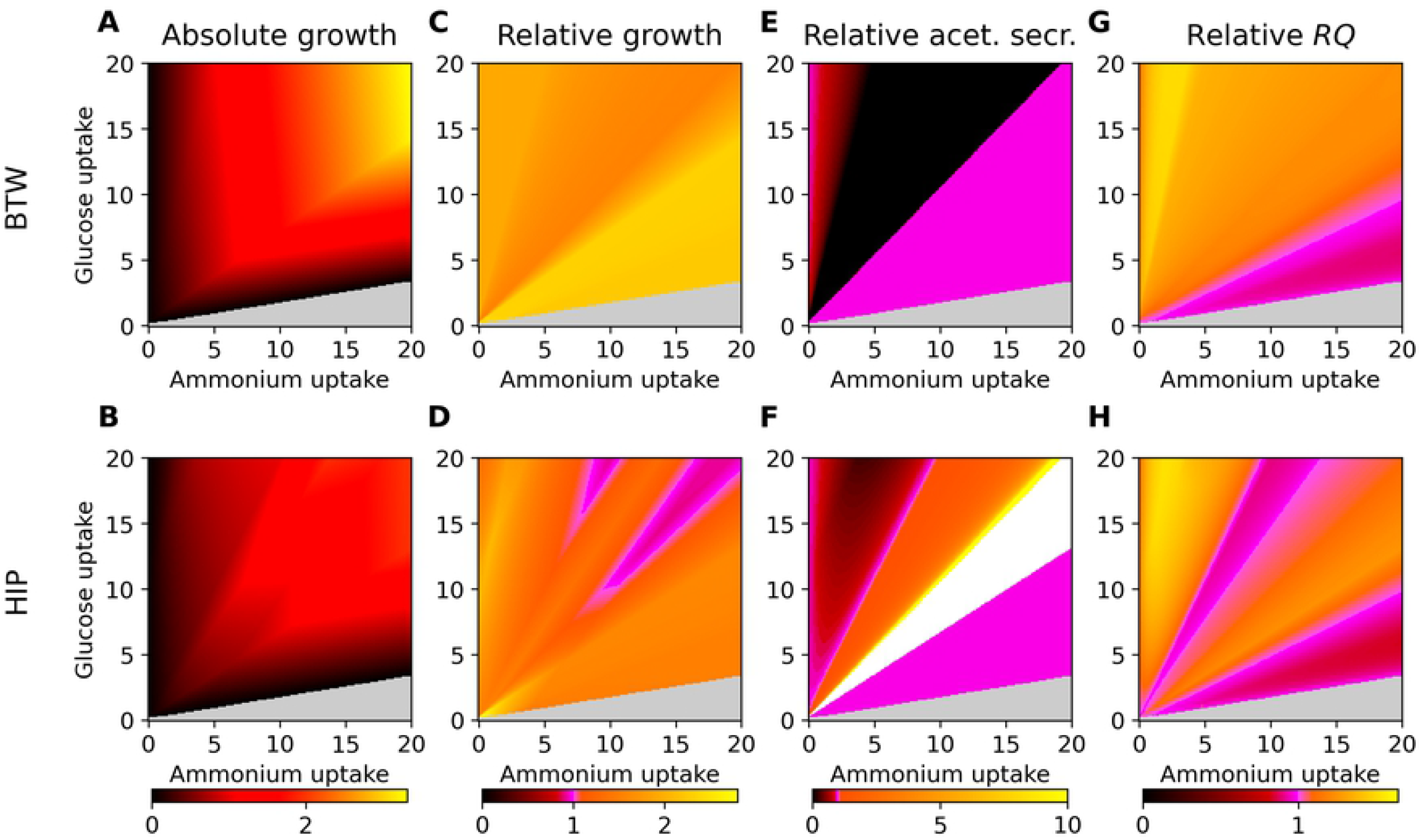
Side-by-side comparison of the BTW and HIP method, panels **C** to **H** are relative to the respective panels (and uptake rates therein) in Fig. 1 of the unlimited environment UL. Glucose and nitrogen were fixed to the given uptake rates, all other uptakes are unconstrained. Non-growth areas are in light grey, a value of unity is indicated by pink. Panels **A** and **B** show growth phenotype phase-planes, panels **C** and **D** show relative growth phenotype phase-planes with the UL BOF as reference. Panels **E** and **F** show acetate secretion flux rates relative to those of the UL BOF, and **G** and **H** show the respiratory quotient *RQ* relative to the UL BOF RQ. The white area in the relative acetate secretion plot for the HIP method indicates a small acetate production in HIP, and zero acetate production in the UL BOF.

We first generated phenotypic phase-planes in response to varying glucose and ammonium uptake rates for the two approaches, as shown in Fig. 3 panels **A** and **B**. If BTW and HIP returned the same results as the UL BOF with standard FBA calculations, the plots in panels **C** through **H** would only be pink. The growth potential is clearly affected by the chosen method: BTW displays higher growth potential compared to the HIP approach. This is to be expected since the BTW methods chooses the combination of available BOFs that maximizes the objective function; in this case total growth. Next, we evaluated the growth potential of the methods relative to the original UL BOF, as seen in Fig. 3 panels **C** and **D** The BTW strategy generates the highest relative growth potential, which is as expected due to its definition. The values are consistently larger than the UL reference BOF. In contrast, HIP generates some areas (shown in pink, panel **D**) with similar growth as the UL BOF. The smallest pink region around coordinate (8.5, 18) is expected, since this is the region in which UL is defined to be the experimental BOF. Similarly, we observe pink at and near this coordinate in both panels **F** and **H**.

The relative *RQ* profiles are shown in Fig. 3 panels **G** and **H**. BTW generates a higher *RQ* than the UL BOF, in the areas with *RQ* < 1, BTW and HIP perform similar to UL. The HIP method therefore generates a BOF biased more towards fermentative states and consequently towards higher acetate secretion rates than BTW does (Fig. 3 panels **E** and **F**). The relative acetate secretion of BTW shows similar phases as the NL BOF. In fact, the absolute acetate secretion, shown in Supplementary S3 Fig.1, indicate a high similarity between the BTW and NL BOFs.

Since fixing uptakes is inconsistent with the definition of HIP-I, we scanned the space of glucose an ammonium uptake fluxes in a different manner. Instead of fixing the uptake fluxes, we enforced an upper bound on their value (and zero as lower bound), at which the HIP-I method was initiated. The resulting growth phenotype phase-plane is shown in Fig. 4 **A**. When using a discrete mapping such as HIP-I, it is to be expected that the queried flux values give rise to two categorically different types of solutions: Either the flux coordinate is a fixed point, for which the HIP-I produces a self-consistent solution without any iteration (i.e. the HIP solution), or it is an unstable point. In the latter case, HIP-I will iterate through a sequence of such points until a fixed point is encountered. Fig. 4 **B** shows one such transition corresponding to the indicated starting and end point in panel **A**.

**Fig 4.**
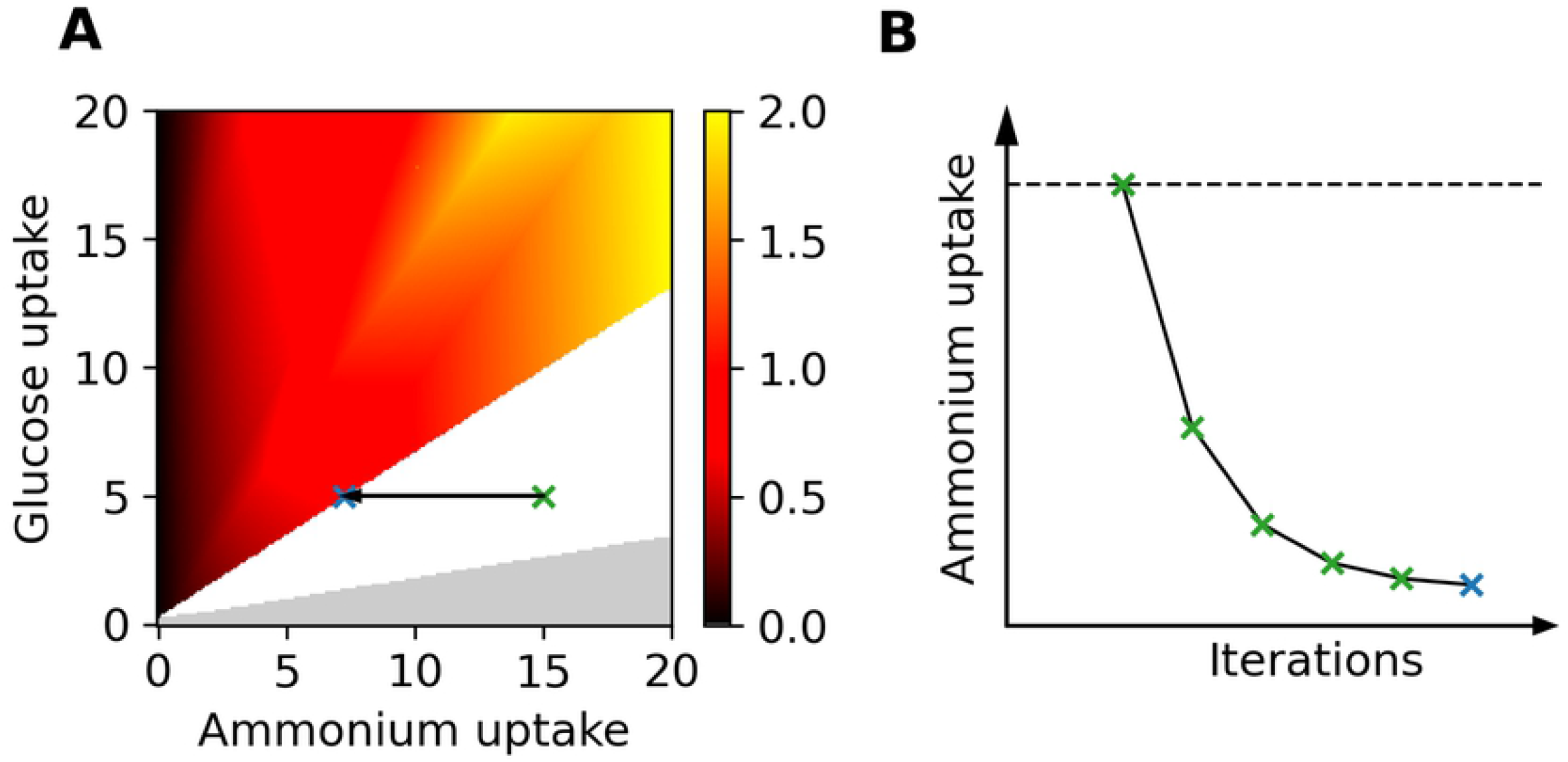
Panel **A** shows the growth phenotype phase-plane of the HIP-I method. A white area indicates no stable HIP-I solution. Gray color indicates a non-growing region in the UL. The remainder of the plot consists of stable HIP-I solutions, i.e. points that have the same growth phenotype as HIP. The dotted arrow between the green and the blue X shows the transition from an unstable HIP-I point to a stable one. Panel **B** illustrates the HIP-I algorithm: The starting constraint indicated by the blue line, here being ammonium uptake, generates a new BOF for the next iteration. This process is repeated iteratively until a stable uptake (within a given tolerance) is found, indicated here by the blue cross.

As expected, a high-ammonium and low-glucose environment is not beneficial for the model and consequently, the uptake is iteratively adjusted to reach a lower nitrogen uptake. Hence, we find that the transition line between the unstable and stable HIP-I coordinates serves as an attractor for the unstable region.

Gene essentiality is an important phenotype in a genome-scale metabolic model [29, 35, 36]. To assess the consequence of the three different BOF formulations and the methods BTW, HIP and HIP-I on gene-essentiality predictions, we conducted a full single-gene knockout screen at 10 different coordinates in the environmental space. For the three different BOF formulations, we used standard FBA calculations to calculate mutant growth rates. Note that, while Fig. 4 shows that HIP-I fixed-point regions show the same growth characteristics (and also other phenotypes) as HIP, their evaluations of knockouts differ. The reason for this is that each mutant genome-scale metabolic network will have a different topology for its fixed-point region. Consequently, a given (C/N) flux coordinate point that is a HIP-I fixed point in the wild-type model, may become HIP-I unstable in a knockout mutant model. Due to the need to run HIP-I through multiple iterations, computational run-times required for a full single-gene knockout screen increase significantly.

The results of the knockout study are shown in Fig. 5, and a depiction of the chosen glucose and ammonium (C/N) coordinates is given in Supplementary S4 Fig.2. A table summarizing the knockout data is to be found in Supplementary S2 Table. When comparing the UL, NL, and CL BOFs for the given environment location, the number of essential genes show small variations in a range from 240 to 250. The effect of gene knockouts in the CL BOF fall into only two categories: Either the gene is essential or it will not affect the growth rate. For the NL BOF, which has the largest carbohydrate fraction, we find the largest fraction of genes with slight to moderate reduction of relative growth, within the relative growth range (0.5, 0.98] in all the selected environments. Additionally, in 7 out of 10 knockout environments, eight genes reduce growth to [0.01, 0.50] of the WT growth rate. In summary, the NL BOF demonstrates more genes with effects on growth than the UL and CL BOFs.

**Fig 5.**
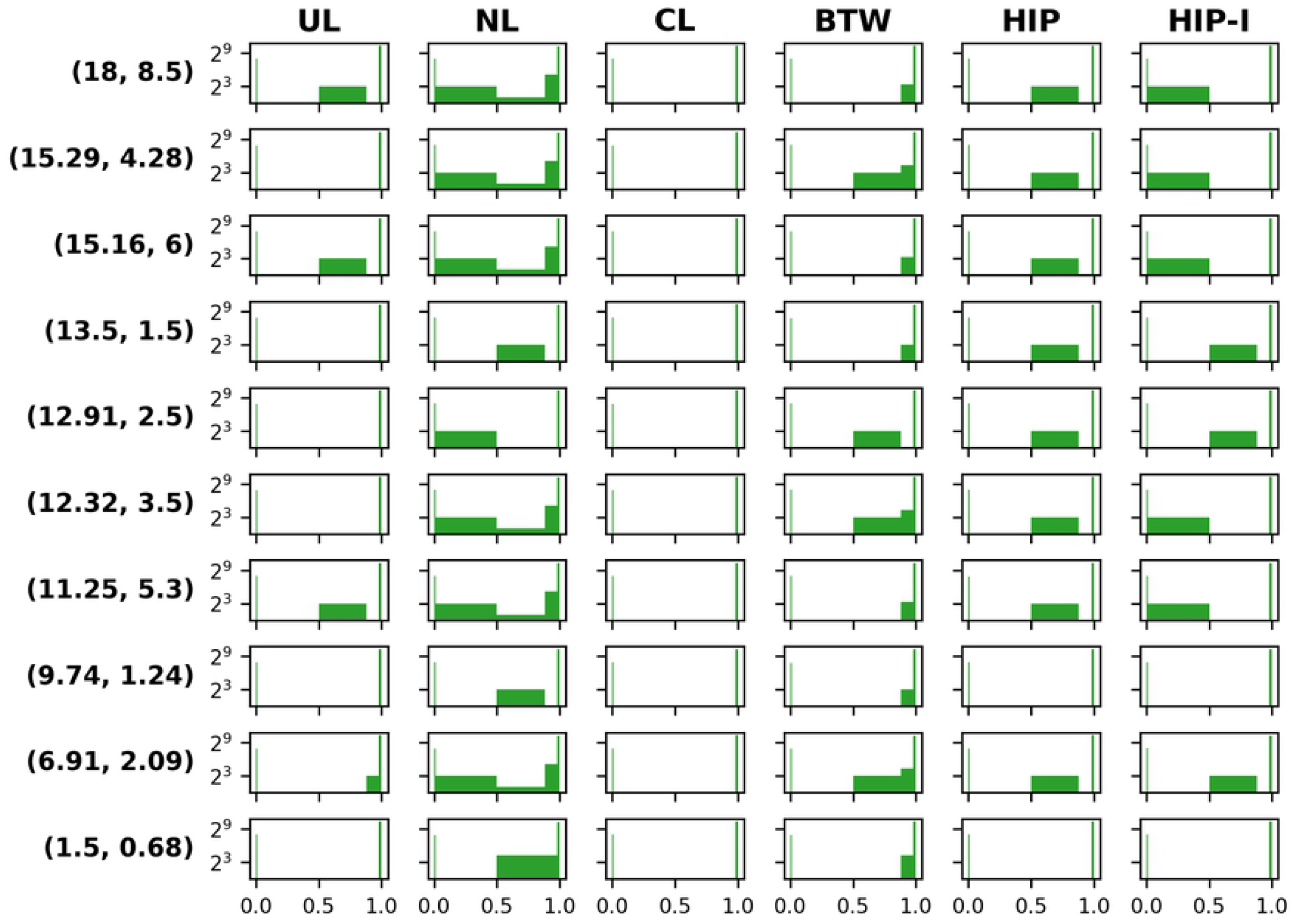
Single gene knockouts at 10 different environments using FBA with the biomass functions being UL, NL, or CL, and for the methods BTW, HIP and HIP-I. The environments are defined by their carbon and nitrogen uptake fluxes, where the flux value represents the maximum uptake limit. The genes are sorted according to mutant growth rate relative to the wild type, sorted into the following intervals: [0, 0.01) / [0.01, 0.50] / (0.50, 0.88] / (0.88, 0.98] / (0.98, 1] These intervals are reflected in the width of each bar in the histograms, while the heights are the log base 2 of the number of genes falling within the respective interval.

Comparing the three methodologies BTW, HIP, and HIP-I, the numbers of essential genes show the largest variation with BTW BOFs: They range from 233 to 250, whereas for HIP we find values from 239 to 250. In 9 out of 10 environments, the model with the HIP-I BOFs contain 250 essential genes. The BTW is the only method where genes with a low relative growth reduction (0.88, 0.98] are found in 9 out of 10 environments. In contrast, the HIP algorithm finds eight genes with intermediate effect in the range (0.5, 0.88] in 8 out of 10 environments, BTW in 4 out of 10 environments. Note that, the number of genes varying in the range of [0.01, 0.88] is same (eight) the for all three methods. These eight genes display a significant reduction of relative growth (down to [0.01, 0.50]) for 5 out of 10 knockouts environments for a HIP-I BOF, similar to the NL BOF. None of the other methods or BOFs contain these genes with a significant reduction. In summary, the BTW BOFs show a larger variation in genes with little phenotypic effects, HIP-I demonstrates more genes with significant growth reductions and more stable number (upper limit) of essential genes.

## Conclusion and Outlook

We implemented three different BOFs in the *i*ML1515 genome-scale metabolic model for *E. coli*. These gave rise to different phenotype profiles for growth potential, acetate secretion potential, gene essentiality, and respiratory quotient. The underlying reason for these changes are metabolic rearrangements necessitated by the different drain on resources imposed by the changes in biomass compositions.

Using this basis, we investigated the challenge of how to implement and combine several different BOFs in a genome-scale model. The approaches presented lead to different formulations of the biomass function, which cause distinctly different phenotypic responses to changes in the nutrient environment. We have proposed two basic methodologies, BTW and HIP, with an iterative extension of the latter in the form of HIP-I. The proposed methods are merely a first line of suggestions to initiate the exploration of this important extension to constraint-based analyses, since high-quality experimental data do not yet exist to evaluate their efficacy. The chosen methods are therefore designed to be fast and easy to apply and interpret, while at the same time having a simple and sensible, appealing heuristic basis. We foresee possible future expansions of these methods or altogether completely different approaches.

Given enough data, one can imagine the development of a range of methods, based on methodologies such as advanced statistical regression analyses, or other machine learning approaches, and mechanistic models involving the specific topology of the metabolic networks. In that respect, non-uniform mapping between nutrient- and biomass space could be considered within the range of perturbed environmental variables. We are aware that there are multiple options to be explored and validated when *in vivo* data become available. The experimental data might not be as linear as we have assumed in this manuscript; non-linear surfaces or step-wise variations are possible consequences of gene-regulatory effects. Additionally, it is not a given that the measurements will fit on a plane due to uncertainties in biomass-component measurements. The HIP and HIP-I methods may easily be extended to this case by generating a best-fit plane using simple linear regression. However, with the presented work, we open a direction for further interdisciplinary discussion on this matter.

Furthermore, it seems reasonable to assume that, for some organisms, the biomass composition could change significantly between quite similar nutrient conditions. An example of this could be a carbon-limiting minimal medium with different carbon sources, such as glucose or lactate. As the biomass composition depends on the environment, detailed and accurate knowledge of environmental conditions is crucial when using multiple BOFs.

We propose that next-generation GSMs should contain (hard-coded) laboratory data about their included BOF(s). Community-driven adjustments to the current version of the *SBML* format might be required, as has recently happened for a different challenge [37]. Including additional fields in the SBML format regarding the BOF offers the possibility of a diverse range of methods for BOF combinations, whether based on the methods presented here, or something completely different. Functions for correct BOF selection, automatic BOF selection based on environmental similarity, or even generation of more suitable BOFs based on a mechanistic understanding of the relationship between environment and biomass composition are all possibilities. We believe that adjustment and combination of BOFs as presented here will also be instrumental in attaining an increased level of mechanistic knowledge of the relationship between environment, metabolism, and biomass composition.

## Materials and methods

In this manuscript, we consider the biomass objective function (BOF) for an organism as follows:

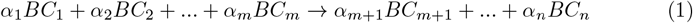

Each compound in the BOF corresponds to a dimension in an *n*-dimensional biomass component space, where the coefficients *α_i_* may vary in response to changes in the organism’s nutrient environment. Note that, in the biomass component space, one particular value for the different *α_i_* corresponds to a single point.

### Generation of artificial biomass functions

In order to test the effects of combining BOFs in multiple ways, different BOFs are needed. The reconstruction *i*ML1515 includes two separate biomass functions with reaction codes 2669 (BIOMASS_Ec_iML1515_core_75p37M) and 2670 (BIOMASS_Ec_iML1515_WT_75p37M). In order to create a single reference biomass, we took the arithmetic mean for each biomass component in the two to generate a new BOF. In Supplementary S1 Table we listed the coefficients *α_i_* of the two BOFs, and the *WT* version contains 31 compounds more than *core*. The new, nutrient-rich BOF is listed as *Mean* and was scaled to 1gh^−1^gCDW^−1^ based on the given molecular weights. We use this BOF as our reference biomass, and we refer to it as the unlimited (UL) environment composition. Note that, the compound *adocbl* (adenosylcobalamin) is present in the WT BOF (BIOMASS_Ec_iML1515_WT_75p37M) but needed to be removed since it inhibits model growth.

Also shown in Supplementary S1 Table is the group we used for each compound. For example, we categorize the compound arg_L[c] arginine in the group *protein* in Tab. 2. Subsequently, all compounds were scaled according to their listed group to generate the nitrogen limited (NL) and carbon limited (CL) environment BOFs. The BOF was scaled so that consumed metabolites for a unit flux through it sums to 1gh^−1^gCDW based on the compounds’ molecular weight in the model.

**Table 2.** The table shows the three environments defined by carbon (C) and nitrogen (N) uptake and the scaling of biomass groups. The groupings are used to scale the containing compounds for the creation of the BOF. For example, we define that protein is scaled by 0.2 in a nitrogen limited environment. Consequently, the factor for each compounds in the protein group is multiplied by the corresponding factor for the new BOF.

| Uptake rates | UL | NL | CL |
| --- | --- | --- | --- |
|  | C<sub>18</sub>, N<sub>8.5</sub> | C<sub>13.5</sub>, N<sub>1.5</sub> | C<sub>1.5</sub>, N<sub>0.68</sub> |
| DNA | 1 | 1.1 | 1.9 |
| RNA | 1 | 0.86 | 1.6 |
| Protein | 1 | 0.2 | 3.5 |
| Lipid | 1 | 20 | 0.26 |
| Carbohydrates | 1 | 15 | 0.2 |
| Energy | 1 | 4 | 0.5 |
| Co-factors | 1 | 4 | 3 |
| Ions | 1 | 1 | 1.1 |
| Others | 1 | 3 | 2 |

**Table 3.** Single gene knockouts at 10 different environments using FBA with the biomass functions being UL, NL, or CL, and for the methods BTW, HIP and HIP-I. The environments are defined by their carbon and nitrogen uptake fluxes, where the flux value represents the maximum uptake limit. Number of genes are reported according to induced mutant growth phenotype relative to the wild type in the following intervals: [0, 0.01) / [0.01, 0.50] / (0.50, 0.88] / (0.88, 0.98] / (0.98, 1].

| (C,N) max flux | UL | NL | CL | BTW | HIP | HIP-I |
| --- | --- | --- | --- | --- | --- | --- |
| 1 : (18, 8.5) | 250/0/8/0/1258 | 250/8/2/35/1221 | 250/0/0/0/1266 | 250/0/0/10/1256 | 250/0/8/0/1258 | 250/8/0/0/1258 |
| 2 : (15.29, 4.28) | 241/0/0/0/1275 | 250/8/2/35/1221 | 241/0/0/0/1275 | 250/0/8/20/1238 | 250/0/8/0/1258 | 250/8/0/0/1258 |
| 3 : (15.16, 6) | 250/0/8/0/1258 | 250/8/2/35/1221 | 241/0/0/0/1275 | 250/0/0/10/1256 | 250/0/8/0/1258 | 250/8/0/0/1258 |
| 4 : (13.5, 1.5) | 241/0/0/0/1275 | 240/0/8/0/1268 | 240/0/0/0/1276 | 233/0/0/8/1275 | 240/0/8/0/1268 | 250/0/8/0/1258 |
| 5 : (12.91, 2.5) | 241/0/0/0/1275 | 250/8/0/0/1258 | 241/0/0/0/1275 | 250/0/8/0/1258 | 250/0/8/0/1258 | 250/0/8/0/1258 |
| 6 : (12.32, 3.5) | 241/0/0/0/1275 | 250/8/2/35/1221 | 241/0/0/0/1275 | 250/0/8/20/1238 | 250/0/8/0/1258 | 250/8/0/0/1258 |
| 7 : (11.25, 5.3) | 250/0/8/0/1258 | 250/8/2/35/1221 | 241/0/0/0/1275 | 250/0/0/10/1256 | 239/0/8/0/1269 | 250/8/0/0/1258 |
| 8 : (9.74, 1.24) | 241/0/0/0/1275 | 240/0/8/0/1268 | 241/0/0/0/1275 | 233/0/0/8/1275 | 240/0/0/0/1276 | 250/0/0/0/1266 |
| 9 : (6.91, 2.09) | 241/0/0/8/1267 | 240/8/2/35/1231 | 241/0/0/0/1275 | 240/0/8/20/1248 | 240/0/8/0/1268 | 250/0/8/0/1258 |
| 10 : (1.5, 0.68) | 247/0/0/0/1269 | 245/0/19/18/1234 | 249/0/0/0/1267 | 235/0/0/19/1262 | 249/0/0/0/1267 | 258/0/0/0/1258 |

Since the existing knowledge of biomass composition as a function of microbial growth environment is somewhat limited, we generated artificial biomass functions for three different environments, all of which are under aerobic conditions in a minimal mineral medium: (1) CL - carbon limitation (glucose), (2) NL - nitrogen limitation (ammonia), and (3) UL - an environment without any nutrient limitation. These three artificial BOFs contain distinct and biologically plausible differences in the amounts of the five major groups of macromolecules: DNA, RNA, proteins, lipids, and carbohydrates. All compounds appear in all environments, and we assume no abrupt change in the presence of a single compound, such as the poly(3-hydroxybutyrate) in *A. latus* [28].

We define measured uptake rates for glucose and ammonium, the other uptake rates required for growth in a minimal medium are set to be non-limiting. Specifically, this refers to the uptake rates of oxygen and the other compounds set to −1000 mmol gCDW^−1^ h^−1^, which is also their default values included in the original genome-scale model. In Tab. 2 the chosen environments are defined by the carbon (C) and nitrogen (N) uptake rates and their scaling of the macromolecular groups in the BOF is presented. Here, we will assume that RNA is 1.6 times higher in a carbon starvation (CL) environment (C_1.5_, N_0.68_), and 0.86 times less abundant in a nitrogen starvation (NL) environment (C_13.5_, N_1.5_), than in the unlimited (UL) environment (C18, N_8.5_). The scaling applies equally to all compounds within a defined group, details of which can be found in Supporting Information S1 Table. The relative placement of the three biomass functions in the considered 2-dimensional (ammonium, glucose) nutrient space is depicted in Supplementary S4 Fig.2.

### Method 1: Biomass Tradeoff Weighting (BTW)

The standard flux balance analysis (FBA) problem can be stated as the following linear program [1]:

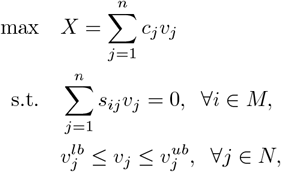

where *S_ij_* is the stoichiometric coefficient of metabolite *i* in reaction *j, v_j_* is the flux through reaction *j*, 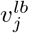 and 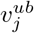 are respectively the lower and upper bounds on reaction *j*. Here, *C_j_* is the objective function coefficient of reaction *j*, typically chosen such that the only non-zero coefficient corresponds to the reaction flux of the BOF.

The BTW method is based on the assumption that, given enough time, bacteria will obtain a biomass composition most optimal for their growth circumstances: If a bacterium in a population shifts its composition slightly to attain a higher growth rate without impeding any other functions or utility, then with time, this adaptation will be propagated by natural selection. As bacteria grow and produce copies of themselves, they use the resources available to the best of their evolved ability. It is therefore possible that, given some limiting nutritional factor on growth, the biomass composition would adapt to use as little as possible of that limiting factor.

While conceptually simple, this method is even simpler in implementation: several biomass compositions for the organism are acquired, then implemented in the standard FBA formulation. All the different biomass functions are included in the objective function by assigning their objective function coefficients *C_j_* = 1, with all other entries being zero. This allows the linear optimization procedure to distribute the optimal flux simultaneously among the multiple BOFs in order to produce the objective value. An illustration of this method for a simple case in 1 dimension is displayed in Fig. 2 **A**. Thus, there is no explicit dependence of the BTW biomass objective function coefficients on the nutrient uptake fluxes.

### Method 2: Higher-dimensional-plane InterPolation (HIP)

This approach is based on assuming the existence of a stable, linear mapping between the nutrient environment space and the biomass coefficient space. This is conditional on similar nutrient environment conditions generating similar values for the organism’s biomass composition. When this assumption holds, it is possible to infer the biomass composition for an organism growing under specified growth conditions if one knows the biomass composition when the organism grows under similar conditions. From this line of thought stems the Higher-dimensional-plane InterPolation (HIP) method.

We hypothesize that the assumption of linearity in the biomass component space will be reasonable if the points in the uptake rate space are close together. We illustrate HIP for a simple case of mapping from one environmental parameter to the amount of one biomass coefficient, see Fig. 2.

In this paper, we have implemented and tested the HIP approach for the hypothetical case of only two nutrient uptake fluxes, those of glucose and ammonium, affecting the biomass composition of the model *i*ML1515 for *Eschericha coli*. In order to uniquely determine the HIP linear mapping between this (2-dimensional) nutrient environment space and the biomass component space, we need to have three points in nutrient environment space (that are not in a line) and their corresponding mappings in biomass component space. For this, we use the UL, CL and NL BOFs and their corresponding location in nutrient environment space.

We implement HIP by using either the upper bounds or fixed values of the nutrient environment uptake fluxes to determine the relevant BOF composition. Subsequently, we perform a standard FBA analysis using this BOF.

#### Higher-dimensional-plane InterPolation-Iteratively (HIP-I)

As the name would imply, this is an iterative extension of the HIP method that originates from the fact that the optimal solution to HIP may consist of nutrient environment space uptake fluxes that are inconsistent with the values used for determining the HIP biomass function: For a HIP simulation mirroring standard FBA, we constrain the problem using upper bounds on the nutrient environment uptake fluxes. The nutrient environment space coordinates corresponding to these upper bounds are used to determine the specific biomass composition in HIP. If the optimal solution has determined values for these nutrient uptake fluxes that are different from the upper bounds, the solution is not self-consistent. In such a case, the new fluxes through the uptake reactions are used as a new input to the HIP function, and the process is then repeated iteratively until a self-consistent solution is determined, as illustrated in Fig. 4 **B**.

We implement HIP-I as follows. First, we define *u* as the vector of upper bounds for the nutrient environment uptake fluxes, and *v_k_* as the optimal flux solution after the k-th iteration of HIP-I. The initial step of HIP-I (*k* = 0) uses *u* to determine the BOF (*B*_0_). Running HIP with these inputs results in an optimal uptake flux vector *v*_1_. If |*v_i_* − *u*| ≤ *ϵ*, the algorithm terminates, and the HIP-I solution for *u* will equal the HIP solution for *u*. Instead, if |*v*_1_ − *u* > *ϵ*, we use *v*_1_ and its corresponding biomass composition *B*_1_ as input for the next (*k* = 1) HIP execution. This is iterated until |*v_k_* − *v*_*k*-1_| ≤ *ϵ* or until we reach *k_m_*, a predetermined maximum number of iterations. For the current simulations, we used *ϵ* = 10^−3^ and *k_m_* = 100. Note that *k_m_* was never reached in any of the iterations, in HIP areas *k_m_* was *k_m_* = 0 and in other areas *k_m_* ≈ 2.5.

### Modeling and plotting

All simulation analyses have been implemented using the COBRA Toolbox 3.0 [38] in Matlab 2019b [39], and we used the genome-scale metabolic reconstruction *i*ML1515 for *Eschericha coli*. The COBRA Toolbox function *optimizeCbModel* was used in combination with the solver *gurobi* [40]. The settings *feasTol* and *optTol* were changed from 10^−6^ to 10^−9^ to resolve numerical issues for all calculations. For knockouts, we used the COBRA toolbox function *singleGeneDeletion* with the data output *grRatio*. The threshold for a knockout was 10^−2^. The *RQ* determinations were performed with parsimonious FBA [41]. The result of the first optimization for the BOF was fixed. Afterwards, the optimized acetate production flux was also fixed. Then, the parsimonious FBA minimizes all fluxes in the (irreversible) model. This allows for stable, minimal required fluxes of oxygen and CO_2_ to determine RQ. The plots were created with Matplotlib [42]. Values are for the most part represented exactly as they are in the colormaps, with the exception that the values for relative acetate are capped at 10, due to some values being divided by very low ones.

## Acknowledgments

EA and CS would like to thank ERA-IB-2 and The Norwegian Research Council grants #271585 and #294605; EA and TK the Norwegian Research Council grant #248885, and EA and EK the Norwegian Research Council grant #269084.

## Declarations

The funders had no role in study design, data collection and analysis, decision to publish, or preparation of the manuscript. The authors have declared that no competing interests exist.

## Supporting information

**S1 Table Three biomasses used and scaling of groups.** This table lists the three BOFs for the nitrogen- and carbon-limitation, as well as the unlimited function. Note, that these BOFs are made up and not based on measurements. Additionally we list the groups used in the manuscript for scaling shown in Tab. 2.

**S2 Table Table of knockouts.**

**S3 Fig.1 Direct plots for the Fig. 3.** The plot shows the non-relative heatmaps presented in Fig. 3, for BTW, HIP and UL for a direct comparison. Note especially the light-grey zero-areas in the direct comparison.

**S4 Fig.2 Selected points for single-gene knockout analysis presented in Fig. 5.** The figure shows the glucose-ammonium environmental space and the three BOFs UL, NL, and CL (marked in red). The other points marked in the figure are the coordinate locations for the single gene knockout analysis. The numbering of the points corresponds to the numbering in Fig. 5.

